# Maximizing immunopeptidomics-based bacterial epitope discovery by multiple search engines and rescoring

**DOI:** 10.1101/2024.11.22.624860

**Authors:** Patrick Willems, Fabien Thery, Laura Van Moortel, Margaux De Meyer, An Staes, Adillah Gul, Lyudmila Kovalchuke, Arthur Declercq, Robbe Devreese, Robbin Bouwmeester, Ralf Gabriels, Lennart Martens, Francis Impens

## Abstract

Mass spectrometry-based discovery of bacterial immunopeptides presented by infected cells allows untargeted discovery of bacterial antigens that can serve as vaccine candidates. However, reliable identification of bacterial epitopes is challenged by their extreme low abundance. Here, we describe an optimized bioinformatical framework to enhance the confident identification of bacterial immunopeptides. Immunopeptidomics data of cell cultures infected with *Listeria monocytogenes* were searched by four different search engines, PEAKS, Comet, Sage and MSFragger, followed by data-driven rescoring with MS^2^Rescore. Compared to individual search engine results, this integrated workflow boosted immunopeptide identification by an average of 27% and led to the high-confidence detection of 18 additional bacterial peptides (+27%) matching 15 different *Listeria* proteins (+36%). Despite the strong agreement between the search engines, a small number of spectra (< 1%) had ambiguous matches to multiple peptides and were excluded to ensure high-confident identifications. Finally, we demonstrate our workflow with sensitive timsTOF SCP data acquisition and find that rescoring, now with inclusion of ion mobility features, identifies 76% more peptides compared to Q Exactive HF acquisition. Together, our results demonstrate how integration of multiple search engine results along with data-driven rescoring maximizes immunopeptide identification, boosting the detection of high-confidence bacterial epitopes for vaccine development.

## INTRODUCTION

Antimicrobial resistance is an increasing worldwide healthcare threat^1^, urging preventive measures that include the development of novel bacterial vaccines^2^. The discovery of bacterial epitopes and antigens is a crucial step to inform on protective vaccine formulations. In adaptive cellular immunity, peptides derived from pathogenic proteins are presented by the major histocompatibility complex (MHC) – human leukocyte antigen (HLA) in human – in order to recognize and clear infected cells. Self-peptides and peptides derived from intracellular pathogens can be presented by MHC class I molecules after their generation via proteasomal degradation and further proteolytic maturation, leading to presentation of typically 9-mer peptides by human HLA molecules^3^. Isolation and mass spectrometry (MS)-based identification of such peptides presented on the surface of infected host cells is possible by so-called immunopeptidomics, a valuable technique for untargeted identification of putative bacterial antigens in the context of vaccine design^4^. In recent years, immunopeptidomics screens have identified epitopes of various bacterial pathogens, including *Listeria monocytogenes*^5^, *Mycobacterium tuberculosis*^6^ and *Salmonella typhimurium*^7^.

Thirty years after the first MS-based detection of a handful of immunopeptides^8^, contemporary MS-based proteomics achieves the sensitive identification of thousands of immunopeptides per sample. Increasing depth in immunopeptides is facilitated by low-input MS instruments such as the timsTOF instruments that incorporate ion mobility (IM) separation^9,10^. TimsTOF instruments apply parallel accumulation-serial fragmentation (PASEF) which accumulates a package of ions while simultaneously separating another package of ions that is serially fragmented upon elution from the IM cell^11,12^. Specifically for immunopeptidomics, customized ion mobility-mass isolation windows, so-called thunder polygons due their characteristic shape, have been applied to include singly charged immunopeptides^9,10^. Instruments not equipped with an IM device are not able to distinguish singly charged peptide ions from singly charged contaminants, and these ions are therefore often excluded. To improve the MS-based detection of immunopeptides, TMT derivatization has been used to promote multiply charged peptide precursors and the abundance of b-ions^13,14^. Next to data acquisition, several advances were made in the computational identification of immunopeptides from the MS data. As immunopeptides contain non-tryptic ends, the complexity of the peptide search space is vastly increased compared to standard tryptic peptide searches. Consequently, immunopeptide identification is more challenging, and different search engines have been shown to exert a certain bias and low consistency between identified immunopeptides^15,16^. However, extending peptide-to-spectrum match (PSM) search engine scores with additional orthogonal scoring features based on predicted fragment ion intensities^17-19^, retention time (RT)^20^ and more recently IM^21,22^, increases the confidence and number of identified peptides. Such data-driven rescoring was shown to boost immunopeptide identification over 30%^9,19,22^ and with the recent TIMS^2^Rescore algorithm up to 71% more MHC class I immunopeptides were reported when leveraging spectrum, RT and IM predictions^23^.

Despite these technological advances, confident identification of presented bacterial peptides remains particularly challenging due their extremely low abundance^4^. In a typical infection experiment over 99% of the identified immunopeptides are host self-peptides. The remaining peptides are bacterial immunopeptides that require thorough inspection. Follow-up experiments to ascertain identified bacterial peptides as true hits are of the essence, especially given that these identifications can be decisive in formulating vaccine constituents, for which experimental testing is time-consuming and costly. Therefore, for further validation of bacterial immunopeptides, synthetic peptides or heavy spike-in peptides are often used, where co-elution and similar MS/MS fragmentation of the bacterial peptide and synthetic counterpart should be observed^5,6^. In addition, within a bacterial infection set-up, inclusion of non-infected host samples can serve as a valuable negative control to select for bacterial immunopeptides^5^. Recent studies on cultured cells infected with *Mycobacterium tuberculosis* and *Listeria monocytogenes*, a foodborne bacterial model pathogen further referred to as *Listeria*, revealed bacterial proteins represented by multiple distinct peptides, thereby underlining their immunogenic potential and allowing their prioritization as vaccine candidates^4,5^. Despite showing huge potential, the data processing of these studies relied on a single search engine without the benefit from recently developed spectral rescoring tools.

To maximize the performance of immunopeptidomics data analysis for bacterial epitope discovery, we here describe a workflow integrating the results of four search engines, PEAKS Studio 12, Comet, Sage and MSFragger, that includes rescoring with MS^2^Rescore for each engine in parallel. Applying this workflow to label-free and TMT-labeled fractions of a previously published immunopeptidomics dataset from *Listeria*-infected cells^5^ resulted in an additional 18 *Listeria* peptides matching 15 proteins, despite more stringent selection criteria that avoid ambiguous spectrum-to-peptide assignments. Moreover, re-injection and data analysis using a timsTOF SCP instrument nearly doubled the amount of identified immunopeptides and allowed the detection of yet unobserved *Listeria* antigen candidates. Taken together, we describe a highly performant computational framework for immunopeptidomics studies and demonstrate its potential for bacterial antigen discovery.

## EXPERIMENTAL PROCEDURES

### System information

PEAKS Studio 12^24^ and IsobaricAnalyzer^25^ were ran on a 20 CPU Windows system (64 GB RAM, 2.75 GHz, AMD EPYC 9454 processors). All other search engines, rescoring, FlashLFQ quantification and automated reports were ran on a 32 CPU Linux Ubuntu (v20.04.06) system (512 GB RAM, 2.75 GHz, AMD EPYC 9454 processors) in an automated end-to-end search-to-reporting Python script.

### Immunopeptide identification

Q Exactive HF (.RAW) and timsTOF SCP (.d) data was searched by four search engines in parallel: (*i*) MSFragger version 4.1^26^, (*ii*) Comet version 2023.01 rev. 2^27^, (*iii*) Sage version 0.14.7^28^, and (*iv*) PEAKS Studio 12 (build 20240709)^24^. The raw mass spectral data was first pre-processed and searched by MSFragger, whereafter recalibrated mzML files^29^ were used for Sage, Comet and PEAKS Studio. In case of PEAKS studio, no mass correction or chimera scan analysis is performed to search an identical set of re-calibrated MS/MS spectra, which facilitates downstream integrations of PSMs. Spectra were searched against a concatenated target-decoy (reverse) database comprised of UniProtKB reference proteomes for human (UP00005640, 20596 proteins) and *Listeria monocytogenes* EGD (UP000016703, 2847 proteins). For label-free searches, no static modifications were set, and variable modifications were Cys cysteinylation, Met oxidation, protein N-terminal acetylation and pyro-Glu formation from peptide N-terminal Asp and Glu. For TMT-labeled fractions, variable Met oxidation and Cys cysteinylation, and static TMT10-plex modification of peptide N-termini and Lys. An unspecific peptide digestion option was used, restricting peptide length to 7-20 and mass 600-5,000 Da. For Q Exactive HF data a 10 ppm precursor mass tolerance was applied for all searches, while using default fragment mass tolerances for Comet (fragment bin tolerance 0.02), MSFragger (20 ppm), and PEAKS Studio 12 (0.02 Da). For timsTOF SCP data, 15 ppm were used as precursor and fragment mass tolerance for all engines (0.015 fragment bin for Comet), except setting a 0.03 Da fragment mass tolerance for PEAKS Studio 12. The optimized fragment mass tolerance for the main search set by MSFragger were used for Sage searches (timsTOF SCP and Q Exactive HF data). In case of Sage, RT and IM model fitting was used for PSM scoring, while the Deep Learning Boost option was enabled for PEAKS Studio 12.

### Immunopeptide rescoring and cross-engine integration

Search results of each search engine were rescored separately using MS^2^Rescore^22,30^, and peptides identified at 1% peptide FDR after rescoring by each search engine were aggregated. MSFragger, Sage and PEAKS outputted each a Percolator input (.pin) file with search engine PSM scoring features. PEAKS Studio 12 was configured to export decoy hits (system settings “detach-service” and “export-decoy” set to true) and all PSMs were exported by setting the minimum required - lg10P value to 0. PSM reports with search engine features were formatted by custom Python scripts to a tab-separated value (TSV) file used as input for MS^2^Rescore v3.1.0 (dev9)^22,30^. For both Q Exactive HF and timsTOF data, DeepLC version 2.2.32^20^ was used to score PSMs by generating features describing the deviations to the predicted peptide RT. In addition, MS^2^PIP Immuno-HCD^22^, TMT and timsTOF^9^ models were used to compare and rescore against predicted b/y-ion intensities for Orbitrap label-free, Orbitrap TMT-labeled and timsTOF SCP label-free data, respectively. Lastly, in case of timsTOF data, peptide collision cross section (CCS) values were predicted by IM2Deep version 0.1.3^23^ and used for rescoring. Mokapot^31^ was ran before and after extending the PSM scoring features using the default linear support vector machine (SVM) model, and peptides and their PSMs were filtered at a 1% peptide FDR. We integrated the results of the four search engines at a spectrum-level, keeping track of PSMs supported by multiple search engines or the rare cases (<1%) where a spectrum was matched to distinct peptides. Peptides having ambiguous PSMs were flagged and excluded for further consideration to err on the side of caution.

### Immunopeptide quantification

In case of label-free Q Exactive HF data, FlashLFQ version 1.2.5^32^ was used to perform label-free quantification. A single FlashLFQ input file containing all combined search engine results was used (requiring filename, RT, precursor change, peptide sequence and mass). Matching-between-runs was enabled with a 5 ppm tolerance, a RT window of 2.5 min, requiring two isotopes and only quantifying the identified peptide precursor charge. In case of TMT10plex labeled data, FileConverter and IsobaricAnalyzer modules part of the OpenMS platform^25^ were used to preprocess Thermo RAW files and subsequently extract reporter ion quantifications, respectively. For HeLa data, *Listeria*-infected channels included 127N, 127C, 128N and 128C, whilst for uninfected controls 129N, 129C, 130N and 130C were used. In case of HCT-116 data, *Listeria*-infected channels included 126, 127N, 127C and 128N, whilst for uninfected controls 129C, 130N, 130C and 131 were used. Similar to Mayer et al.^5^, quantification results were filtered for at least two intensity values in either uninfected or *Listeria*-infected conditions in Perseus^33^. Missing values were imputed from a normal distribution around the detection limit.

### HLA binding prediction and Gibbs clustering

Standalone versions of NetMHCpan-4.1b^34^ and NetMHCIIpan-4.3^35^ were used to predict the binding strength of immunopeptides to HeLa and HCT-116 MHC class I and II alleles, respectively. In case of MHC class I, these included HLA-A*03:19, HLA-A*68:02, HLA-B*15:03 and HLA-C*12:03 for HeLa, while HLA-A*01:01, HLA-A*02:01, HLA-B*45:01, HLA-B*18:01, HLA-C*07:01 and HLA-C*05:01 were used for HCT-116. Alleles with the strongest predicted binding (lowest %rank score) were used in the final reports. Unsupervised alignment and clustering of all 8- to 12-mer peptides was performed using GibbsCluster 2.0^36^ using recommended settings for MHC class I immunopeptides of length 8-13 (motif length 9, max deletion length 4, max insertion length 1).

### Reports and visualization

All data processing steps were executed using a custom Python script (Python version 3.8.10). Data organization was handled using pandas^37^. Plots were generated using matplotlib^38^ and seaborn^39^. Sequence logos were generated using Logomaker^40^.

### TimsTOF LC-MS/MS analysis

Immunopeptides from *Listeria*-infected or uninfected HeLa cells were isolated as described^5^. Leftover, dried peptide mixtures eptides re-dissolved in 17 µl loading solvent A (0.1% TFA in water/acetonitrile (ACN) (99.5:0.5, v/v)) of which 8 µl was injected for LC-MS/MS analysis on an Ultimate 3000 RSLC nanoLC in-line connected to a timsTOF SCP mass spectrometer (Bruker). Given that for the previous Q Exactive HF analysis 5 out of 15 µl was used^5^, we injected here approximately 25% of the peptide material compared to the initial study. Trapping was performed at 20 μl/min for 2 min in loading solvent A on a 5 mm trapping column (Thermo scientific, Pepmap, 300 μm internal diameter (I.D.), 5 μm beads). The sample was further separated on a reverse-phase column (Aurora elite 75µm x 150 mm 1.7um particles, IonOpticks) following elution from the trapping column by a linear gradient starting at 0.5% MS solvent B (0.1% FA in water/ACN 20:80 (v/v)) at a flow rate of 250 nl/min for 30 min reaching 37.5% MS solvent B, increasing MS solvent B to 55% MS solvent B after 38 min, finally increasing further to 70% MS solvent B after 40 min, followed by a wash for 5 min and re-equilibration with 99.5% MS solvent A (0.1% FA in water). The flow rate was decreased from 250 nl/min to 100 nl/min at 20 min and increased again to 250 nl/min at 40 min. A ten PASEF/MSMS scan acquisition method was used in DDA-PASEF mode with a precursor signals intensity threshold at 500 arbitrary units. An adapted polygon in the *m/z*-IM plane was used to include HLA-I singly charged precursors (Supplementary Table 1). The mass spectrometer was operated in non-sensitive mode with an accumulation and ramp time of 100 ms, analyzing in MS from 100 to 1,700 m/z. Precursors were isolated with a 2 Th window below m/z 700 and 3 Th above and actively excluded for 0.4 min when reaching a target intensity threshold of 20,000 arbitrary units. A range from 100 to 1,700 m/z and 0.7 to 1.25 Vs cm^-^² was covered with a collision energy applied according to the ion mobility range in Supplementary Table 2.

## RESULTS AND DISCUSSION

### Multiple search engines and rescoring boost immunopeptide identification

In our previous work we identified 68 *Listeria* immunopeptides in infected HCT-116 and HeLa cell cultures using Peaks Studio 10.5 at a 1% PSM Q-value^5^. Purified immunopeptides from four infected and four uninfected replicate cultures were subjected twice to Orbitrap liquid chromatography-tandem mass spectrometry (LC-MS/MS) analysis, once label-free and once in combination with TMT-labeling to increase immunopeptide identification (Figure 1A)^5,13,14^. Here, we aimed to improve the bioinformatical analysis of these data by developing an *in silico* workflow that leverages the complementary power of four search engines with follow-up rescoring of PSMs. Label-free and TMT-labeled runs of both cell cultures were re-searched with MSFragger^26^, Comet^27^, Sage^28^, and PEAKS Studio 12^24^ to identify MHC Class I immunopeptides (see Experimental Procedures). Resulting peptide-to-spectrum matches (PSMs) outputted by each engine were rescored separately using MS^2^Rescore v3^22,30^, extending search engine score features (e.g. hyperscore, mass accuracy, and others) with orthogonal scoring features derived from MS^2^PIP^17^ and DeepLC^20^.

**Figure 1.**
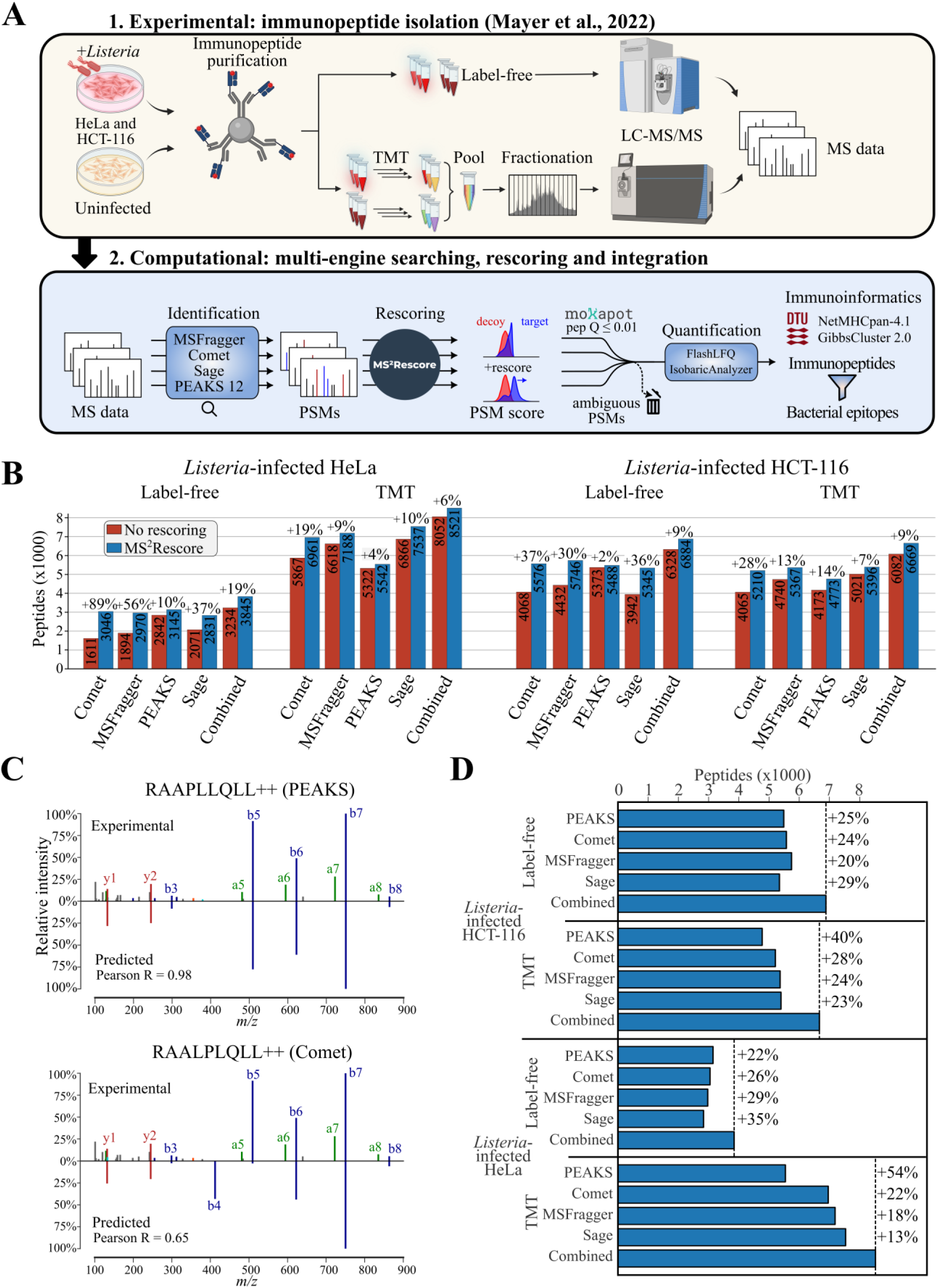
Integration of multiple search engines and rescoring boost immunopeptide identification. (**A**) *Listeria*-infected and uninfected HeLa or HCT-116 cells were subjected to both label-free LC-MS/MS on a Q Exactive HF instrument as well as TMT-labeling and LC-MS/MS analysis on a Fusion Lumos instrument after pre-fractionation^5^. Resulting MS data was searched by four search engines in parallel and the search results of each engine were rescored by MS2Rescore v3^30^ independently. All peptide and PSM identifications with a peptide q-value < 1% were aggregated and further subjected to quantification, HLA binding prediction and Gibbs clustering. (**B**) Rescoring by MS^2^Rescore boosts the number of identified peptide sequences per search engine and across engines. (**C**) Annotated MS2 spectrum (HeLa label-free replicate 2, scan 52812) matched to both ‘RAAPLLQLL’ by PEAKS Studio 12 and ‘RAALPLQLL’ by Comet. The experimental spectrum is displayed on top and the MS^2^PIP-predicted spectrum at the bottom, indicating the Pearson spectrum correlation calculated by MS^2^Rescore (‘spec_pearson’ feature). (**D**) Integrating the results of four search engines after rescoring further boosts immunopeptide detection. The dotted line indicates the total number of unique, identified peptide sequences after rescoring per sample.

Rescoring of the label-free search engine results of Sage, Comet and MSFragger by MS^2^Rescore resulted in a dramatic increase of identified peptide sequences (peptide Q-value ≤ 1%) ranging from 30 up to 89% in the *Listeria*-infected HeLa and HCT-116 cells (Figure 1B). In case of PEAKS Studio 12 the ‘Deep learning boost’ within PEAKS Studio 12 already leverages spectrum and RT scoring features, explaining the high identification numbers prior MS^2^Rescore. Rescoring of TMT-labeled samples also yielded an additional 4 to 28% of identifications in both cell lines. Unlike the Immuno-HCD MS^2^PIP model^22^ for label-free (non-tryptic) immunopeptides, rescoring of TMT-labeled fractions was performed using the MS^2^PIP TMT Orbitrap HCD model. Despite being trained on TMT-labeled tryptic peptides, this model delivered a median correlation between 0.83 – 0.87 (Figure S1). This matches the performance as observed for tryptic peptides^30^, justifying its usage for TMT-labeled immunopeptides.

After rescoring, we integrated the identified peptides and PSMs of the four search engines. A total of 120,696 PSMs (peptide Q-value ≤ 1%) to a single peptide were made by a single or by multiple engines (Data S1A), of which 103,639 (85.9%) were supported by at least two search engines. Next to spectra allocated to a single peptide, 1,010 MS2 spectra (< 1% of total PSMs) were assigned by search engines to different peptides (Data S1B). These mostly included ambiguous matches to peptides with isobaric Ile to Leu substitutions and minor peptide sequence variations. For instance, a spectrum in the HeLa label-free sample was matched to ‘RAAPLLQLL’ by PEAKS and ‘RAALPLQLL’ by Comet – thus peptides with an exchange of the fourth and fifth amino acid (‘PL’ to ‘LP’) that lack fragment ions to distinguish between both peptides (Figure 1C). Such ambiguity due to incomplete fragmentation spectra is prevalent in immunopeptidomics^41^, facilitated by the abundance of candidate PSMs associated with non-tryptic, unspecific searches (especially when searching multiple species). Although scoring features such as the Pearson correlation score, e.g. 0.98 for ‘RAAPLLQLL’ compared to 0.68 for ‘RAALPLQLL’, can suggest the most probable PSM candidate, or chimeric spectra can contain fragments of co-eluting peptides, we err on the side of caution by not considering these PSMs.

At the peptide level, integrating PSMs assigned to a single peptide by one or multiple search engines after rescoring boosted the number of detected immunopeptides on average by 27% (Figure 1D). Only peptides with at least a single uniquely assigned PSM were retained, while keeping track of the number of ambiguous PSMs to flag potentially unreliable identifications. In total there were 20,664 peptides identified with uniquely assigned PSMs (Data S2A). Conversely, 361 peptides that were solely identified by ambiguous PSMs were discarded (Data S2B). As such, the integration of multiple search engine result not only yields additional immunopeptide identifications, but also flags spectrum-to-peptide ambiguities.

After identification, the ensemble of retained PSMs across all search engines is quantified by FlashLFQ^32^ or IsobaricAnalyzer^25^ for label-free and TMT-labeled peptides, respectively. Peptides identified by all four search engines showed the highest intensity, while peptides identified by a single search engine or those gained by rescoring were relatively lower abundant (Figure S2), suggesting that the use of multiple search engines and rescoring benefits the identification of low intensity immunopeptides. In addition, HLA class I binding prediction was performed by NetMHCpan-4.1^34^ and unsupervised alignment and clustering of all 8- to 12-mer peptides was performed using GibbsCluster 2.0^36^. As a first sign of high-quality immunopeptide detection, presented HLA class I peptides were predominantly 9-mers as indicated by the sequence length histogram of identified peptides (Figure 2A). We quantified a total of 9,924 9-mers, which is an additional 1,408 9-mers (+17.5%) compared to our previous study^5^ using PEAKS Studio 10.5 (Figure S3). Moreover, 86 to 91% of identified 9-mers were predicted to exert strong binding (SB) to HeLa or HCT-116 HLA class I alleles (Figure 2A). Evaluating the overlap of identified SB peptides (8 to 12-mers) between the different search engines revealed that 57 to 69% of SB peptides were confidently identified by all four engines (Figure 2B). Conversely, 9 to 13% of all SB peptides were only identified by a single search engine, showcasing how hundreds of immunopeptide identifications are gained by our multi-engine search strategy with follow-up rescoring. As also reflected by the Gibbs clustering analysis (Figure S4-S6), HLA-A*68:02, HLA-B*15:03 and HLA-C*12:03 were predominant alleles in HeLa samples, and HLA*A01:01, HLA*A02:01, HLA*B18:01 and HLA*B45:01 were in HCT-116 cells (Figure 2C). Notably, HLA-B*18:01 presented 245 and 405 SB 8-mer peptides in HCT-116 label-free and TMT samples (Figure S7), respectively, matching the earlier observed peptide length binding preference of this allele^42^.

**Figure 2.**
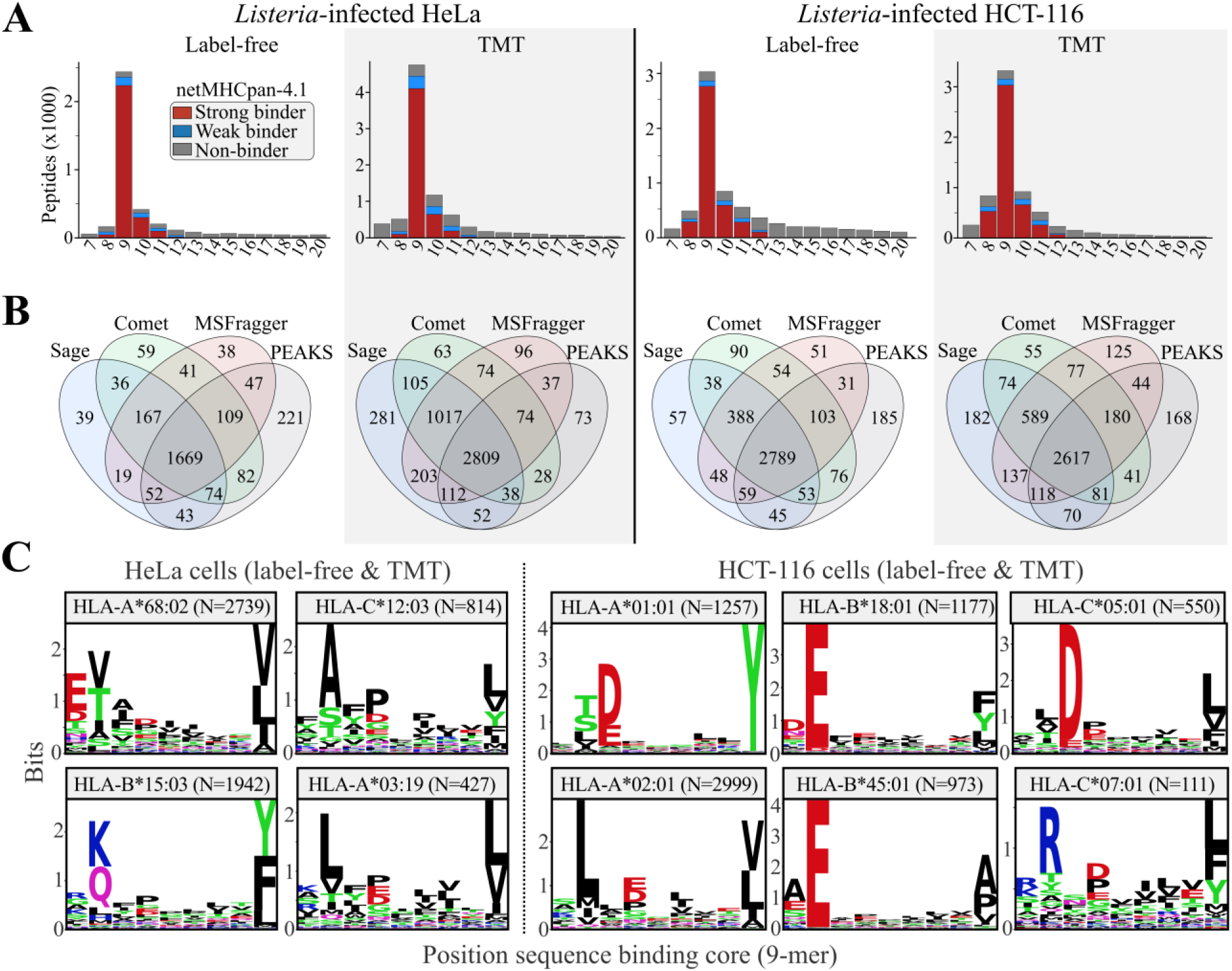
Integration of multiple search engines and rescoring yields complementary immunopeptide identifications. (**A**) The number of unique peptide sequences identified per amino acid length. Peptide sequences predicted as strong binder (SB, %rank < 0.5) or weak binder (WB, %rank < 2) by NetMHCpan-4.1^34^ were indicated in red and blue, respectively. Other peptides (non-binder, NB) were indicated in grey. (**B**) Venn diagram showing the overlap of identified peptide sequences predicted as NetMHCpan-4.1 strong binders per search engine. (**C**) Sequence logo of peptides predicted as strong binder (SB, %rank < 0.5) by NetMHCpan-4.1^34^. Logos were made by Logomaker^40^ using the peptide 9-mer binding core predicted by NetMHCpan-4.1.

### Detection of additional high confident *Listeria* immunopeptides

Within the used host-pathogen infection setup, a major goal is the identification of bacterial *Listeria* immunopeptides. With the bacterial *Listeria* proteome being ∼13-fold smaller than the human proteome and typically far less abundant than the host proteome in infected cells, confident identification of bacterial immunopeptides is challenging. Given that the identified bacterial immunopeptides inform on putative vaccine targets, thorough scrutinization of obtained matches is of the essence. Therefore, multiple quality checks are in place to verify bacterial spectral matches within our workflow (Figure 3A). Of the 173 peptides matched to *Listeria* peptides, 20 *Listeria* peptides were identified from spectra assigned to multiple peptides and therefore removed (Data S3). For instance, in the HCT-116 cells a single spectrum was matched to both ‘TMT-TIDELAGK_TMT_I’ (*Listeria* LMON_0362) by Sage and ‘TMT-TIDEIQK_TMT_L’ (human DDB1) by PEAKS, MSFragger and Comet (Figure 3B). Both peptides are isobaric, with two Leu/Ile substitutions and the combination of Ala and Gly equaling the mass of Gln. The human peptide is most likely true, given its identification by three different search engines and a high Pearson correlation (0.97) to the MS^2^PIP predicted spectrum. Several ambiguous PSMs included matches to peptides with Leu/Ile substitutions, and an additional protein BLAST search of the remaining 153 *Listeria* peptide sequences to an extended human protein database comprised of all human UniProtKB and non-canonical translation proteins of nuORFdb^43^ revealed one additional ambiguous assigned peptide that was also removed (Data S3).

**Figure 3.**
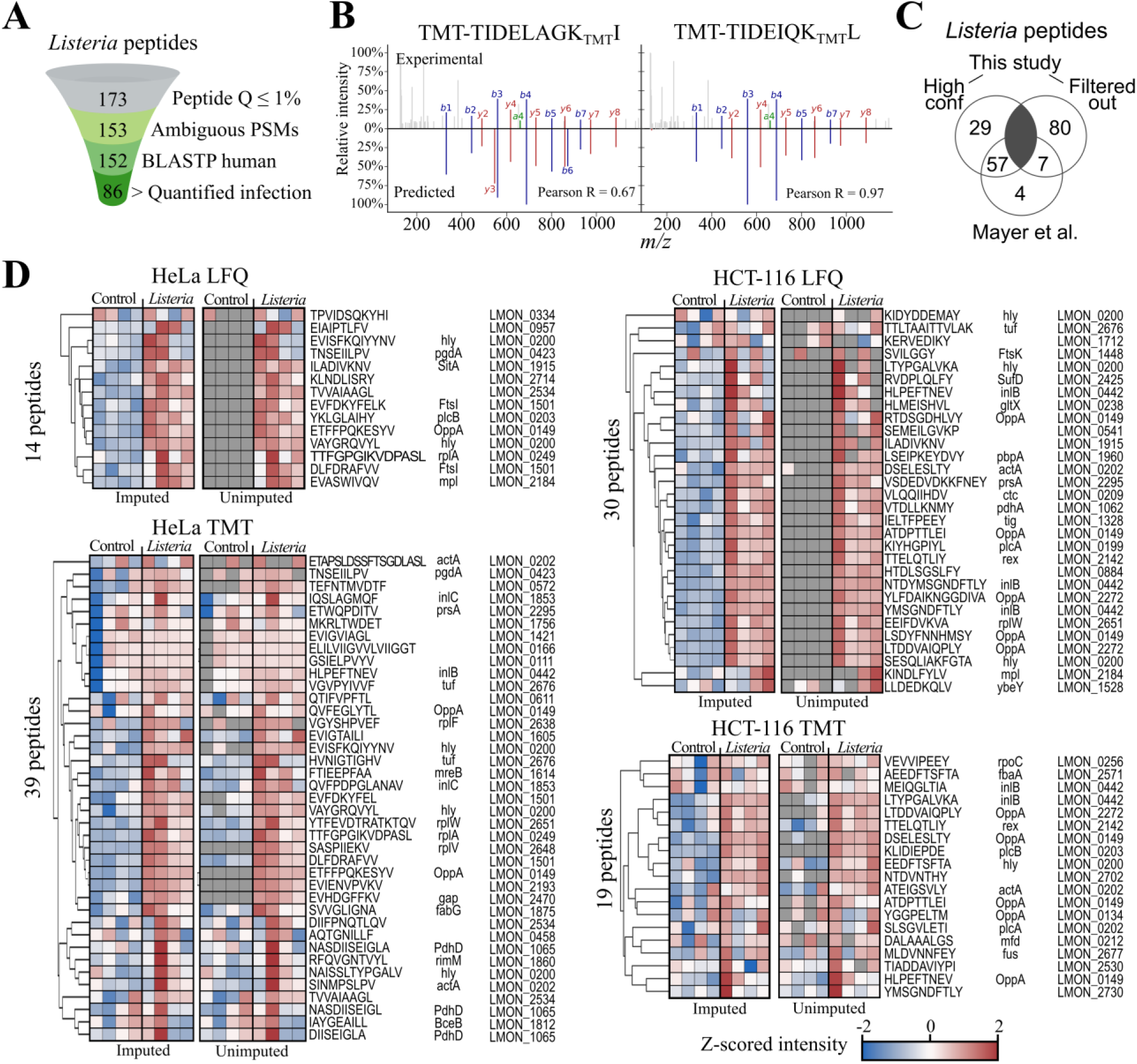
Detection of additional high confident *Listeria* immunopeptides. (**A**) Filters applied for selecting bacterial immunopeptides, resulting in a final number of 86 high-confident *Listeria* peptides. (**B**) Annotated MS2 spectrum (HCT-116 cells TMT fraction 1, scan 19403) matched to ‘TMT-TIDELAGK_TMT_I’ by Sage or matched to ‘TMT-TIDEIQK_TMT_L’ by MSFragger, Comet, and PEAKS. The experimental spectrum is displayed on top and the MS^2^PIP-predicted spectrum at the bottom with their respective Pearson correlation. (**C**) Venn diagram showing the overlap of high-confidence and filtered out *Listeria* peptides in this study to the high-confidence peptides described in Mayer et al.^5^. (**D**) Imputed and unimputed intensity heatmaps of high confident *Listeria* immunopeptides for all four experimental conditions. Z-scored log_2_ intensities were displayed after FlashLFQ^32^ label-free quantification (LFQ) and TMT reporter ion intensities outputted by IsobaricAnalyzer^25^. Missing values were imputed by Perseus^33^.

As a final quality criterium we made use of the uninfected samples as negative control, which logically should lack bacterial peptide identifications or matched intensity signals. Following the filtering criteria of Mayer et al.^5^, we required the remaining 152 *Listeria* peptides to be quantified in at least two out of four *Listeria-*infected replicate samples and with a higher average abundance compared to the uninfected conditions. This resulted in a final number of 86 unique high confident *Listeria* immunopeptides matching 57 proteins (Figure 2D, Data S3), with 69 (80.2%) peptides predicted to bind to HCT-116 or HeLa HLA alleles (NetMHCpan-4.1 %rank ≤ 2) (Data S3). This presents an additional 18 *Listeria* immunopeptides (+26.5%) and 15 proteins (+35.7%) compared to our initial study^5^. While we re-identified 57 of the 68 initially reported peptides (Figure 3C), we identified 29 novel high confident *Listeria* peptides matching to 25 novel antigens, including 23 HLA class I-binding peptides. Conversely, 11 peptides from our previous study were not retained by the present pipeline, of which seven did not meet the set filtering criteria for quantification (Figure 3C). Other peptides such as ‘EVERPSLGV’ were identified at a PSM q-value < 1% after rescoring, but were withheld in our analysis as it had a peptide q-value of 2.1%. Interestingly, although *Listeria* immunopeptides only constitute a small fraction of the identified immunopeptidome, in the label-free datasets these peptides were of similar intensity compared to human immunopeptides (Figure S2D).

### *Listeria* virulence factors are represented by multiple immunopeptides

Similar to our initial study^5^, there is an unequal distribution of the number of identified high confident immunopeptides across the 57 *Listeria* proteins (Figure 4A). Fifteen *Listeria* proteins (previously 13) were represented by multiple immunopeptides (Figure 4A), including five out of six virulence factors encoded in the *Listeria* Pathogenicity Island 1^44^ (Figure 4B): *plcA* (LMON_0199, two peptides), *hly*/LLO (LMON_0200, six peptides), *mpl* (LMON_0201, two peptides), *actA* (LMON_0202, three peptides) and *plcB* (LMON_0203, two peptides). Instead of a single peptide, we now identified two peptides mapping the phospholipase C *plcB*. Also, instead of seven, there were now ten bacterial proteins identified in both infected HeLa and HCT-116 cell cultures, including the rather poorly characterized oligopeptide ABC transporter, periplasmic oligopeptide-binding protein OppA (LMON_0149, seven peptides), a putative cell membrane protein that was again detected as most represented antigen and vaccine candidate^5^. Taken together, these data demonstrate the enhanced capability of our optimized immunopeptide identification and filtering strategy to detect high confident immunopeptides from bacterial origin.

**Figure 4.**
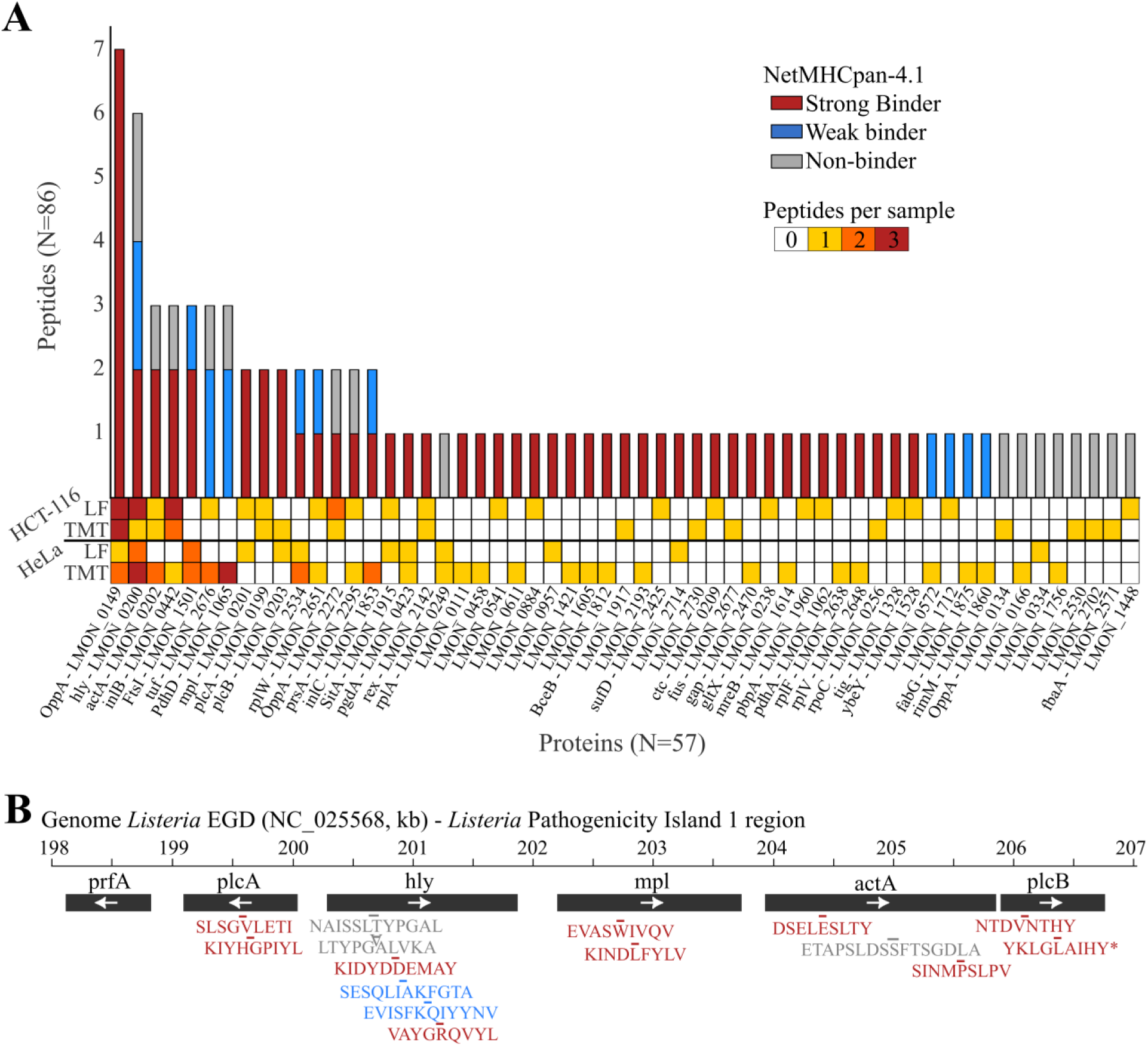
*Listeria* virulence factors are represented by multiple immunopeptides. (**A**) The number of unique immunopeptides identified per *Listeria* protein is shown in a histogram. The number of identified peptides per sample is displayed in a heatmap. (**B**) Genome view of the *Listeria* Pathogenicity Island 1 region^44^ (NC_025568 sequence, numbers in kilobases) with identified immunopeptides for five out of six genes. The plcB peptide ‘YKLGLAIHY’ was a novel immunopeptide identified in this study (*). (**A-B**) Immunopeptides predicted as strong binders by NetMHCpan-4.1 (%rank < 0.5) are indicated in red, weak binders (%rank <2) in blue and non-binders (%rank > 2 or no 8- to 12-mers) in grey.

### Improved immunopeptide identification by DDA-PASEF acquisition

TimsTOF instruments were reported to show enhanced sensitivity in immunopeptide detection^9,10^. To complement the Q Exactive HF-based analysis of our samples reported before^5^ and above, we reinjected 25% of the leftover HeLa label-free samples on a timsTOF SCP instrument. More specifically, we analyzed the samples using 1-hour nanoLC runs coupled to DDA-PASEF data acquisition with an enlarged polygon in the ion cloud plane to include singly charged peptide ions for fragmentation selection, as reported before^9,10^. Again, we integrated the rescored results of four search engines, however, now using the recent TIMS^2^Rescore algorithm that incorporates IM2Deep for prediction and rescoring of peptide collision cross section (CCS) values^23^. Overall, MS^2^PIP-derived features contributed greatly to rescoring (Figure S8), with a median Pearson correlation coefficient of 0.88 was observed for experimental and MS^2^PIP-predicted spectra for PSMs by MSFragger (Figure 5A), corroborating previous timsTOF results^9^. Similarly, we found a strong correlation between predicted and observed RT and CCS values (Figure 5B-C). Rescoring and integration of the four search engine results again significantly boosted the number of identified immunopeptides, delivering a total of 6,876 identified immunopeptide sequences (Figure 5D, Data S4). Similar as other This represents a 79% increase to the 3,845 peptide sequences initially identified on a Q Exactive HF instrument using longer 3-hour nanoLC runs (Data S2A, Figure 1B). 3,363 peptides (out of 6,876 or 49%) were 9-mers of which 2,850 (out of 3,363 or 84%) were predicted as SB peptides by NetMHCpan-4.1 (Figure S9A), again indicating high quality enrichment of immunopeptides in these samples. As anticipated, the enlarged isolation polygon promoted singly charged immunopeptide identifications, with 2,024 9-mer peptides (out of 3,363 or 60%) matched by a singly charged peptide precursor (Data S3). Concomitantly, MHC class I immunopeptides show typical singly charged ion clouds at 850 to 1100 *m/z* and at higher inverse reduced ion mobility (1/K_0_) of ∼1.4-1.6, while doubly charged 9-mers peak at a *m/z* values between 450 to 600 and at ∼0.75-0.9 1/K_0_ (Figure 5E). As anticipated, singly charged peptides were less frequently detected on the Q Exactive HF instrument, with only 23% of 9-mer peptides matched to a singly charged precursor in this dataset (Data S2E). Accordingly, from 1,691 9-mers uniquely detected on the timsTOF SCP but not the Q Exactive HF, 700 were exclusively identified by a singly charged precursor. Looking for bacterial peptides, we identified a total of 48 *Listeria* peptides with unambiguously assigned PSMs and not matching to an extended human database (with Leu/Ile permutation). Since FlashLFQ did not yet support timsTOF label-free quantitation at the time of writing, we continued with 33 *Listeria* peptides that were identified only in *Listeria*-infected replicates but absent in all uninfected replicates, which is more than doubles the amount of *Listeria* immunopeptides detected on Q Exactive HF (14 immunopeptides, Figure 3D). These 33 immunopeptides were derived from 29 *Listeria* proteins (Figure S9B), including eight proteins that were not identified previously in the (re-)analysis of the *Listeria* dataset (Figure 4 and Figure S8). Interestingly, these include iap (invasion-associated protein) or endopeptidase p60, a well-known virulence factor that was shown to be highly immunogenic and facilitate protective immunity against *Listeria*^45,46^.

**Figure 5.**
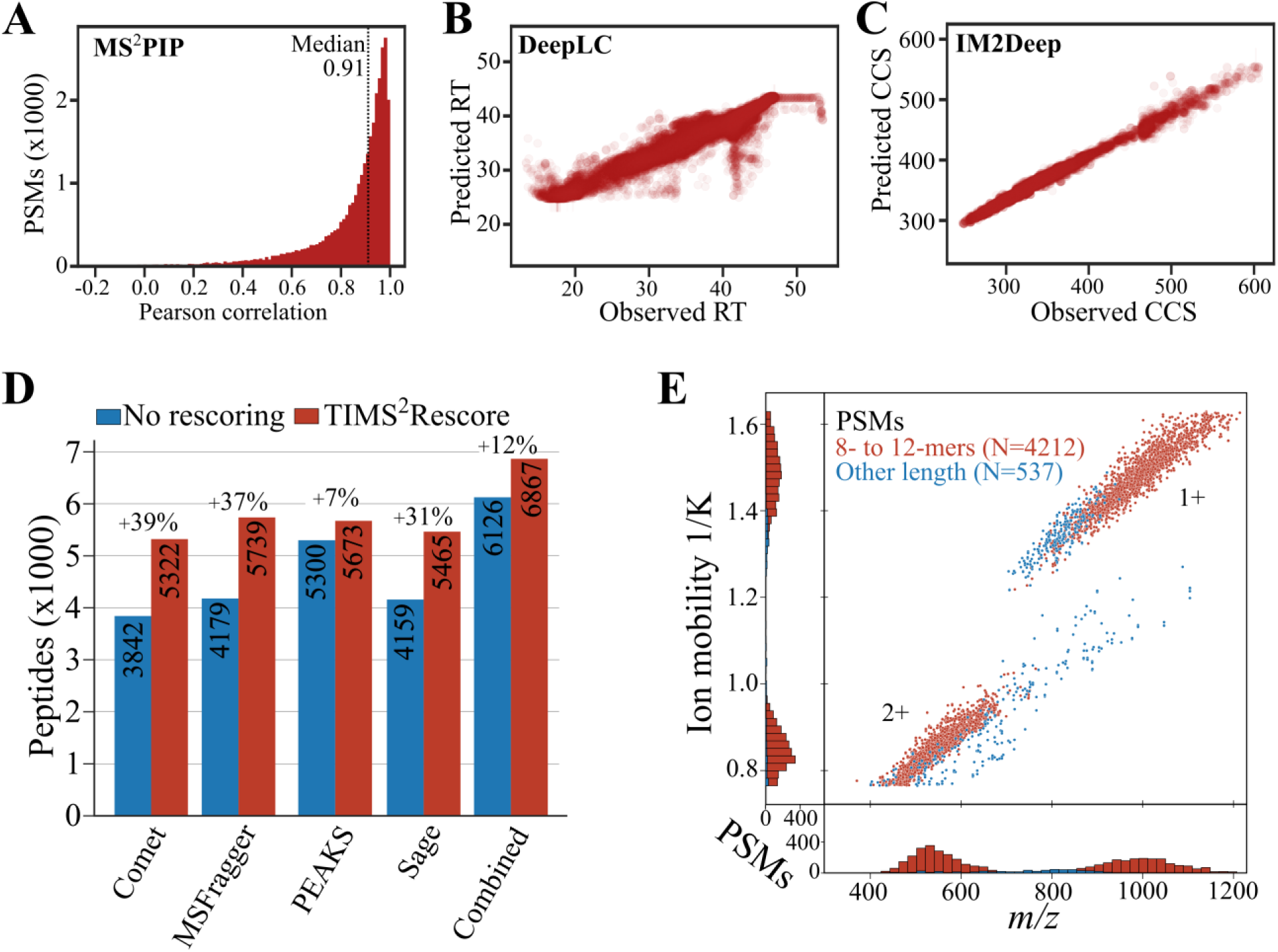
Improved immunopeptide identification by DDA-PASEF acquisition. (**A**) MS^2^PIP timsTOF model performance, displaying a histogram of the Pearson correlation for all PSMs. (**B**) Correlation of DeepLC-predicted and experimental retention time (RT). (**C**) Correlation of IM2Deep-predicted and experimental collision cross section (CCS). (**D**) TIMS^2^Rescore^23^ rescoring and integration of multi-search engine results boosts immunopeptide identification. (**E**) Representative peptide ion identifications plotted across the inversed ion mobility (1/K_0_) and *m/z* dimensions. Identified 9-mer peptides were plotted in red, peptides of other length in blue.

## CONCLUSIONS

We describe a bioinformatical pipeline incorporating four search engines and rescoring to maximize the number of confidently identified bacterial immunopeptides. Overall, we found that immunopeptide identification numbers are boosted both by rescoring and the integration of multiple search engine results. We demonstrate the efficacy of TMT and Immuno-HCD models for rescoring immunopeptides recorded on Q Exactive HF as well as on timsTOF SCP instruments. On the latter instruments, the additional ion mobility dimension was also exploited for rescoring using the novel predictor IM2Deep within TIMS^2^Rescore^23^. Compared to a Q Exactive HF, acquisition on a timsTOF SCP instrument resulted in an overall 76% increase of identified immunopeptides while the number of detected bacterial immunopeptides doubled. Compared to our previous study where 42 *Listeria* antigens were identified, an additional 33 *Listeria* proteins were detected in this work, eight of which were only recorded on the timsTOF SCP (Figure S10). Within this comparison, it should be kept in mind that now a more stringent peptide Q-value of 1% was applied and that PSMs with ambiguous matches to multiple peptides were filtered out. Such additional checking of spectrum-to-peptide ambiguity is important, as ion series are often incomplete and a high ambiguity between host and bacterial peptide precursors is evident due to non-enzymatic search settings. Finally, we envision extension of our multi-engine rescoring pipeline with other search engines compatible with PSM rescoring to further maximize immunopeptide identifications, also for applications beyond bacterial antigen discovery such as viral or cancer (neo)epitope discovery.

## ASSOCIATED CONTENT

### Data availability statement

The timsTOF SCP mass spectrometry proteomics data have been deposited to the ProteomeXchange Consortium (http://proteomecentral.proteomexchange.org) via the PRIDE partner repository^47^ with the dataset identifier PXD055547. Q Exactive HF data from Mayer et al.^5^, re-analyzed in this study, is available via the dataset identified PXD031451. Code used for the automated processing is available at https://github.com/patrick-willems/immunopeptidomics and pipeline output used for this work is deposited at the Open Science Framework (OSF) project with DOI 10.17605/OSF.IO/7DKSA.

### Supporting Information

Supplementary Figure S1. MS^2^PIP model performance for label-free and TMT-labeled immunopeptides per search engine.

Supplementary Figure S2. Peptide label-free intensities per search engine support, rescoring effect and species.

Supplementary Figure S3. Comparison of quantified peptides to our previous study.

Supplementary Figure S4. GibbsCluster 2.0 of HeLa label-free and TMT identified peptides.

Supplementary Figure S5. GibbsCluster 2.0 of HCT-116 label-free identified peptides.

Supplementary Figure S6. GibbsCluster 2.0 of HCT-116 TMT identified peptides.

Supplementary Figure S7. HLA-B*18:01 shows binding preference to 8- and 9-mers.

Supplementary Figure S8. TIMS^2^Rescore feature weights in MSFragger rescoring for MHC class immunopeptides in HeLa label-free samples.

Supplementary Figure S9. TimsTOF identifies novel antigens in *Listeria*-infected HeLa samples.

Supplementary Figure S10. Novel *Listeria* proteins identified in this study.

Supplementary Table 1. Set polygon vertex points for timsTOF SCP re-injection of *Listeria*-infected HeLa label-free samples.

Supplemental Table 2. Set collision energy scheme for timsTOF SCP re-injection of *Listeria*-infected HeLa label-free samples.

Data S1 (XLSX): Peptide-to-spectrum match (PSM) overview of Orbitrap data re-analysis (all samples).

Data S2 (XLSX): Peptide overview of Orbitrap data re-analysis (all samples).

Data S3 (XLSX): Overview of high-confidence and filtered *Listeria* peptides of Orbitrap data re-analysis (all samples).

Data S4 (XLSX): Peptide overview of novel timsTOF SCP analysis for HeLa label-free sample.

## AUTHOR INFORMATION

### Corresponding Author

*Patrick Willems; Francis Impens

## Author Contributions

The manuscript was written through contributions of all authors. All authors have given approval to the final version of the manuscript.

### Notes

The authors declare no competing financial interest.

## ACKNOWLEDGMENTS

We thank the VIB Proteomics Core for LC-MS/MS analysis. P.W., F.T., A.D., R.D., R.B., R.G., L.M. and F.I. acknowledge funding from the Research Foundation Flanders (FWO) [12T1722N, 12AN524N, 1SE3724N, 1SH9O24N, 12A6L24N, 12B7123N, G010023N, G028821N, G0F8616N]. A.G. is supported by a PhD fellowship from the Higher Education Commission (HEC) Pakistan. F.I. and L.M acknowledge support from Ghent University Concerted Research Action grant BOF21/GOA/033 and the Horizon Europe Project BAXERNA 2.0 (101080544).

## ABBREVIATIONS

CCS: collision cross section
LF: label-free
NB: non-binder
SB: strong binder
TMT: tandem mass tag
WB: weak binder

---

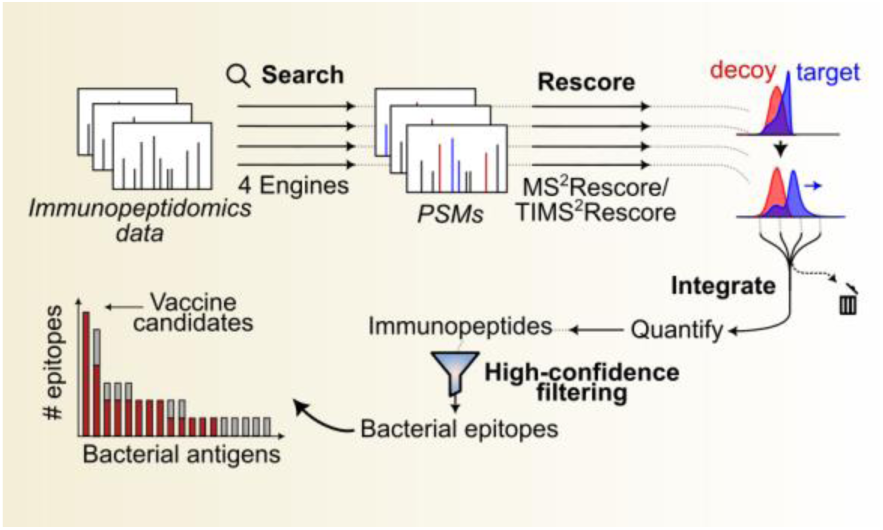
For TOC Only

