## Supplemental Information for "Maximizing immunopeptidomics-based bacterial epitope discovery by multiple search engines and rescoring"

### TABLE OF CONTENTS

|  |  |
| --- | --- |
| Supplementary Figure S1 | MS<sup>2</sup>PIP model performance for label-free and TMT-labeled immunopeptides per search engine. |
| Supplementary Figure S2 | Peptide label-free intensities per search engine support, rescoring effect and species. |
| Supplementary Figure S3 | Comparison of quantified peptides to our previous study. |
| Supplementary Figure S4 | GibbsCluster 2.0 of HeLa label-free and TMT identified peptides. |
| Supplementary Figure S5 | GibbsCluster 2.0 of HCT-116 label-free identified peptides. |
| Supplementary Figure S6 | GibbsCluster 2.0 of HCT-116 TMT identified peptides. |
| Supplementary Figure S7 | HLA-B*18:01 shows binding preference to 8- and 9-mers. |
| Supplementary Figure S8 | TIMS<sup>2</sup>Rescore feature weights in MSFragger rescoring for MHC class immunopeptides in HeLa label-free samples. |
| Supplementary Figure S9 | TimsTOF identifies novel antigens in *Listeria*-infected HeLa samples. |
| Supplementary Figure S10 | Novel *Listeria* proteins identified in this study. |
| Supplementary Table 1 | Set polygon vertex points for timsTOF SCP re-injection of *Listeria*-infected HeLa label-free samples. |
| Supplementary Table 2 | Set collision energy scheme for timsTOF SCP re-injection of *Listeria*-infected HeLa label-free samples. |
| Data S1 (XLSX) | Peptide-to-spectrum match (PSM) overview of Orbitrap data re-analysis (all samples). |
| Data S2 (XLSX) | Peptide overview of Orbitrap data re-analysis (all samples). |
| Data S3 (XLSX) | Overview of high-confidence and filtered *Listeria* peptides of Orbitrap data re-analysis (all samples). |
| Data S4 (XLSX) | Peptide overview of novel timsTOF SCP analysis for HeLa label-free sample. |

**A** HeLa LF samples: Immuno-HCD model

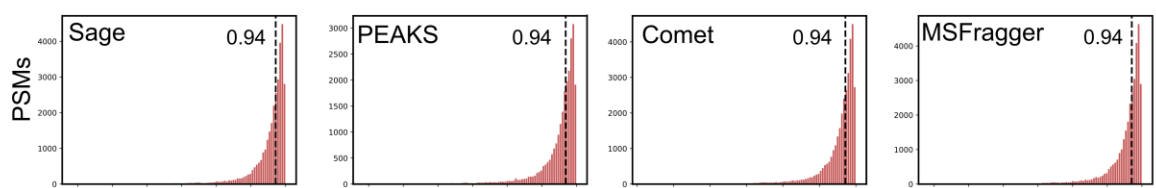

**B** HeLa TMT samples: TMT HCD model

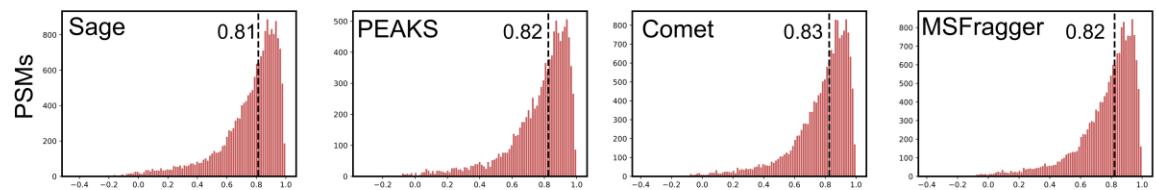

**C** HCT-116 LF samples: Immuno-HCD model

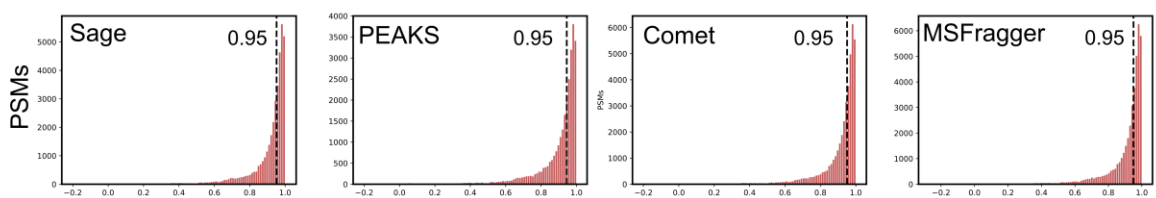

**D** HCT-116 TMT samples: TMT HCD model

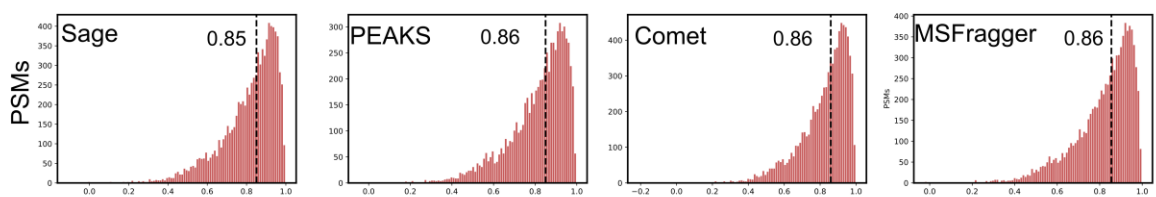

**Supplementary Figure S1. MS<sup>2</sup>PIP model performance for label-free and TMT-labeled immunopeptides per search engine.** TMT10-plex labeled and label-free (LF) immunopeptides were rescored using the MS<sup>2</sup>PIP TMT HCD and Immuno-HCD MS<sup>2</sup>PIP models, respectively, by MS<sup>2</sup>Rescore<sup>1,2</sup>. Pearson correlation histograms of peptide-to-spectrum matches (PSMs) to predicted MS2 spectra were shown for HeLa LF and TMT samples (**A-B**) and HCT-116 LF and TMT samples (**C-D**). The spectral Pearson correlation ('spec\_pearson' feature) was plotted for all target PSMs within 1% mokapot q-value (PSM and peptide level) for all four search engines (*Left to right*). Vertical dotted lines indicate the median Pearson correlation. Abbreviations: LF, label-free; TMT, tandem mass tag.

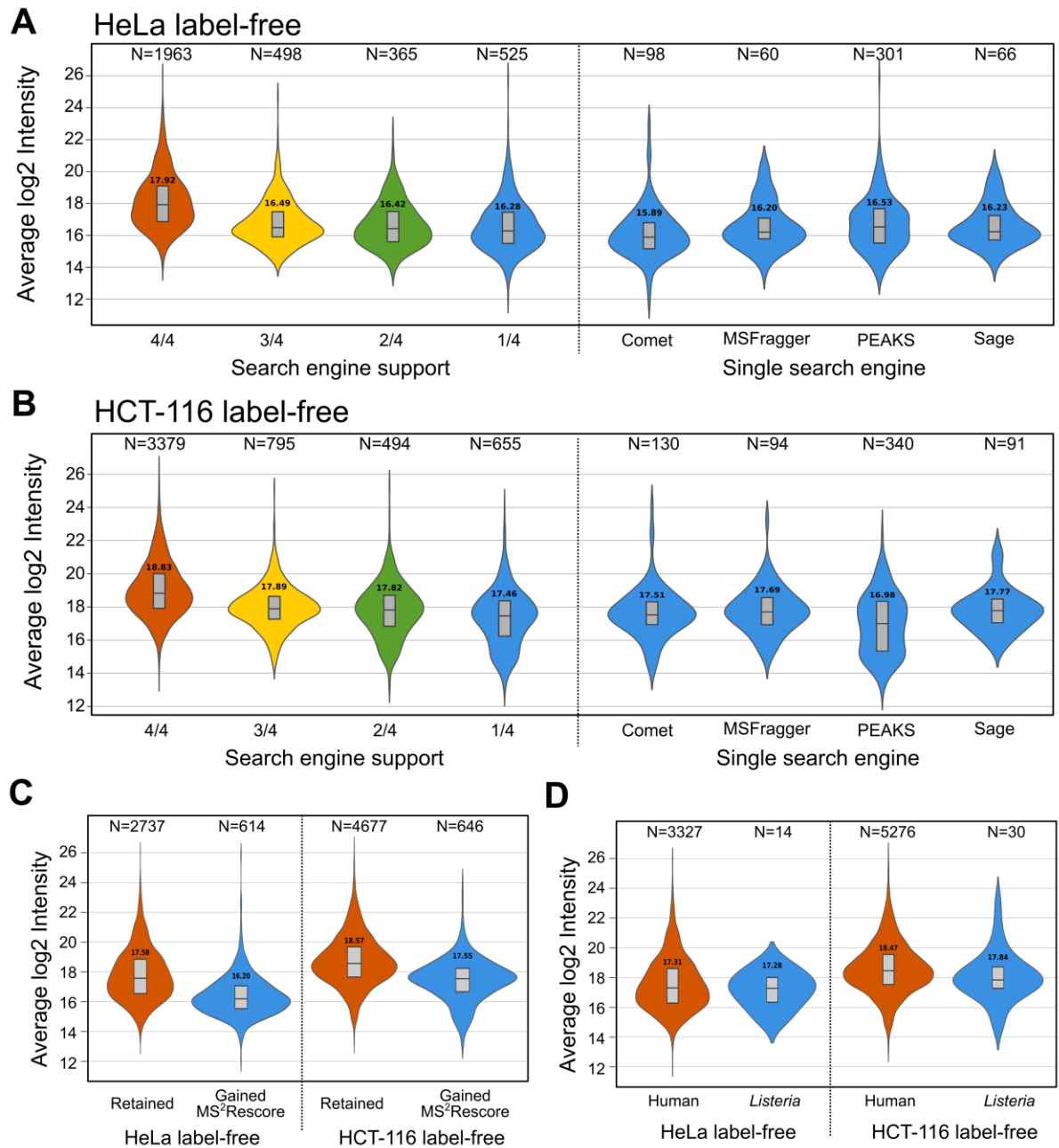

**Supplementary Figure S2. Peptide label-free intensities per search engine support, rescoring effect and species.** Violin plots of the FlashLFQ<sup>3</sup> log<sub>2</sub> intensities (with matching-between-runs) is shown (A-B) according peptide search engine support for HeLa and HCT-116 cells, (C) peptides retained or gained after rescoring by MS<sup>2</sup>Rescore<sup>1,2</sup> (mokapot peptide FDR 1%) and (D) *Listeria* and human self-peptides. Only MHC class I length peptides (8–12 AA) were plotted, except including all high-confidence *Listeria* peptides for the comparison with human self-peptides (D). The number of peptides plotted was indicated above the violin plot, while the median log<sub>2</sub> intensity is indicated above the boxplot inside the violin plot.

### Quantified 9-mers

Recovered: 7815/8444 (92.6%)
Additional: 9924 vs 8444 (+17.5%)

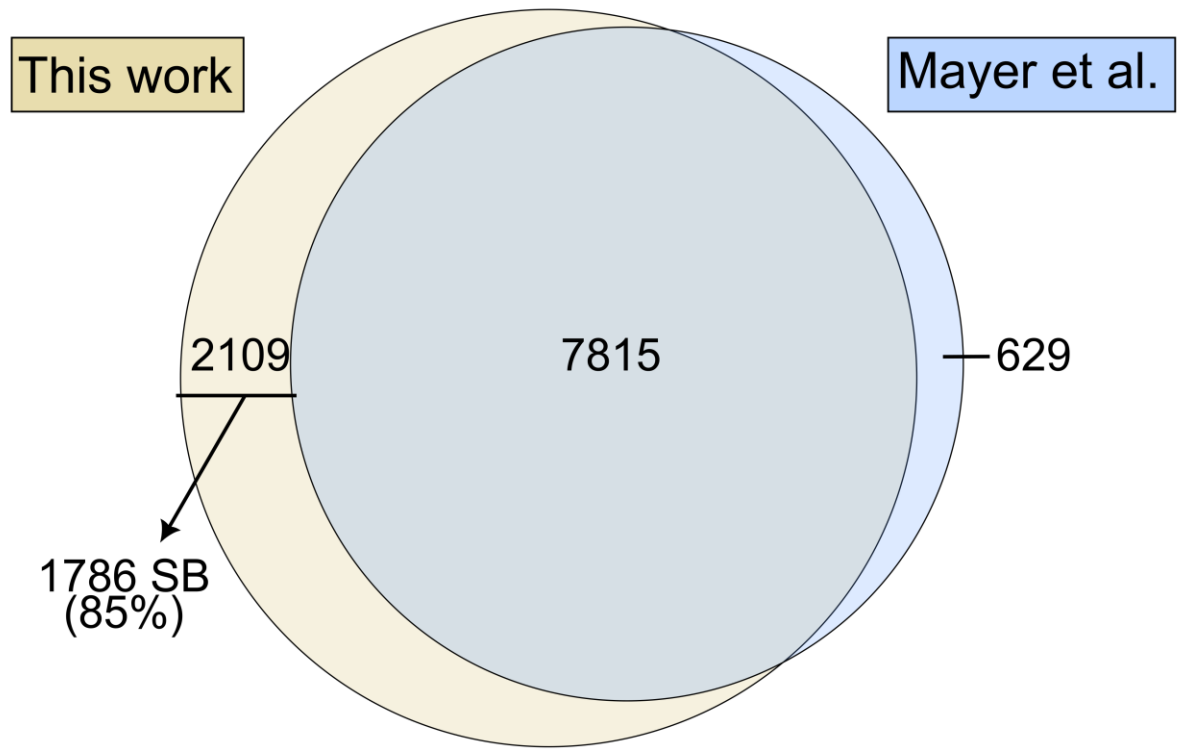

**Supplementary Figure S3. Comparison of quantified peptides to our previous study.** Venn diagram overlap of quantified peptides in this study (9994 peptides) to those by Mayer et al<sup>4</sup> (8444 peptides). Quantified peptides contained at least two valid intensities in either *Listeria*-infected or uninfected samples. Of the 2109 peptides quantified exclusively in this study, 1786 were predicted to exert strong binding to HLA alleles by NetMHCpan-4.1<sup>5</sup>.

### A HeLa - label free

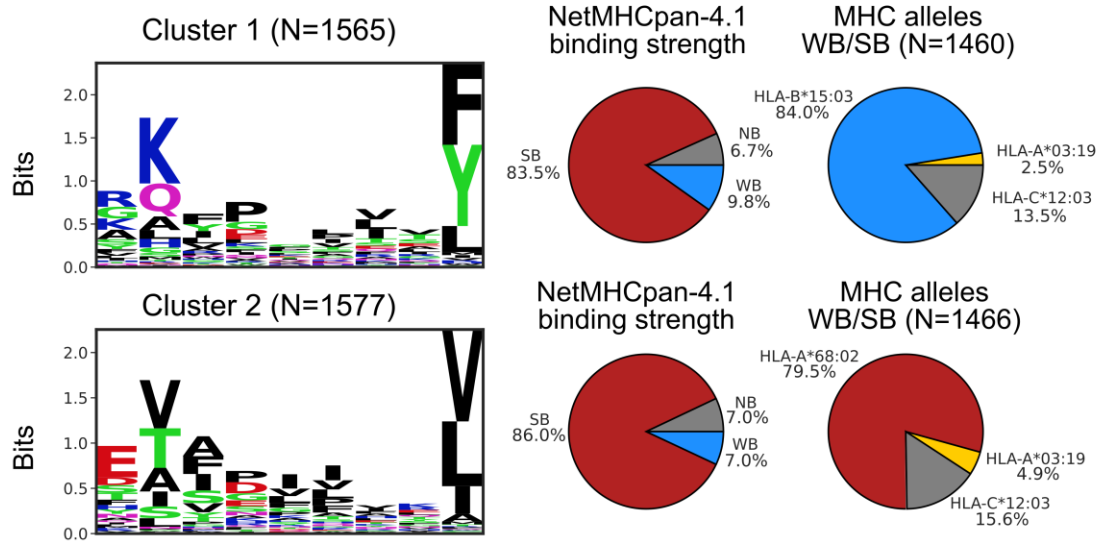

### B HeLa - TMT

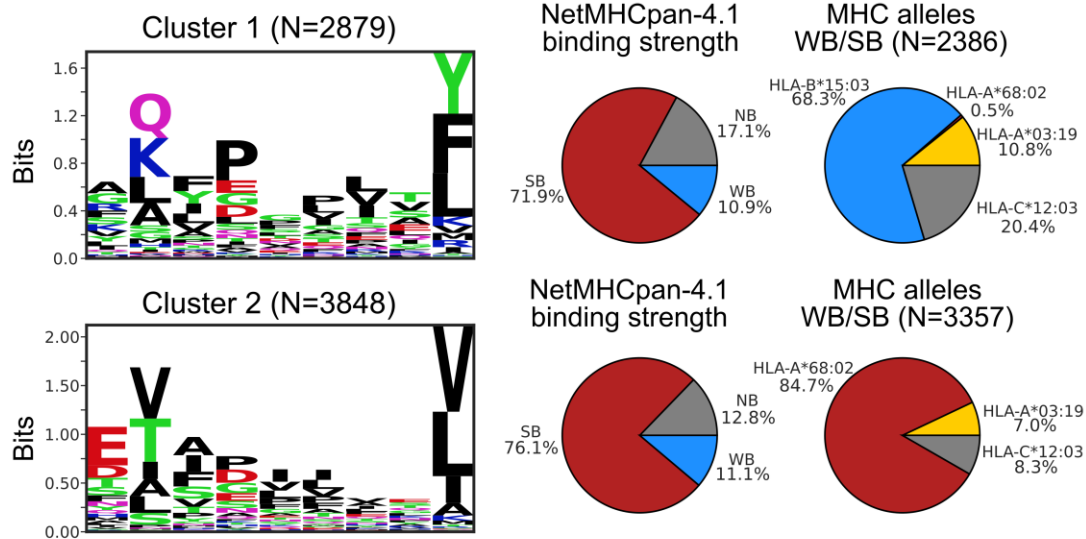

### C NetMHCpan-4.1 naturally presented ligands (EL)

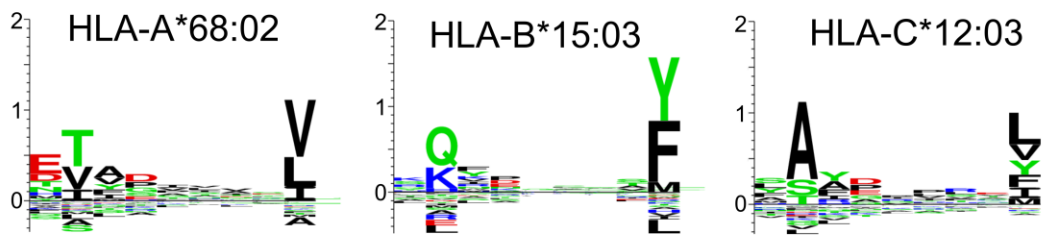

**Supplementary Figure S4. GibbsCluster 2.0 of HeLa label-free and TMT identified peptides. (A-B)** Identified peptides of length 8 to 12 were subjected to unsupervised alignment and clustering by GibbsCluster 2.0<sup>6</sup> using recommended settings. For each cluster, sequence logos were made using Logomaker<sup>7</sup> (*Left*) and the distribution of NetMHCpan-4.1<sup>5</sup> predicted binding level was displayed (*Middle*). For predicted weak and strong binders the HLA allele distribution was displayed (*Right*). (**C**) NetMHCpan-4.1 motif viewer of HeLa alleles (naturally presented ligands, EL). Abbreviations: NB, non-binder; SB, strong binder; WB, weak binder.

### A HCT-116 - label free

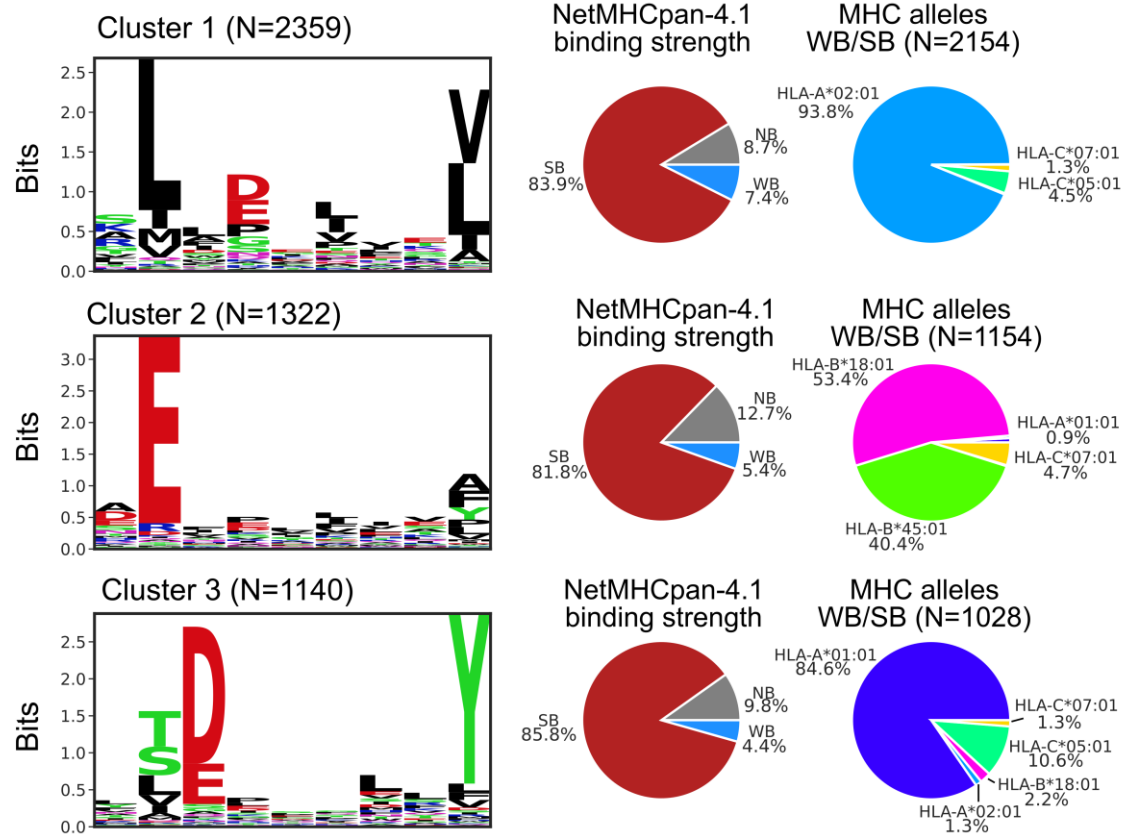

### B NetMHCpan-4.1 naturally presented ligands (EL)

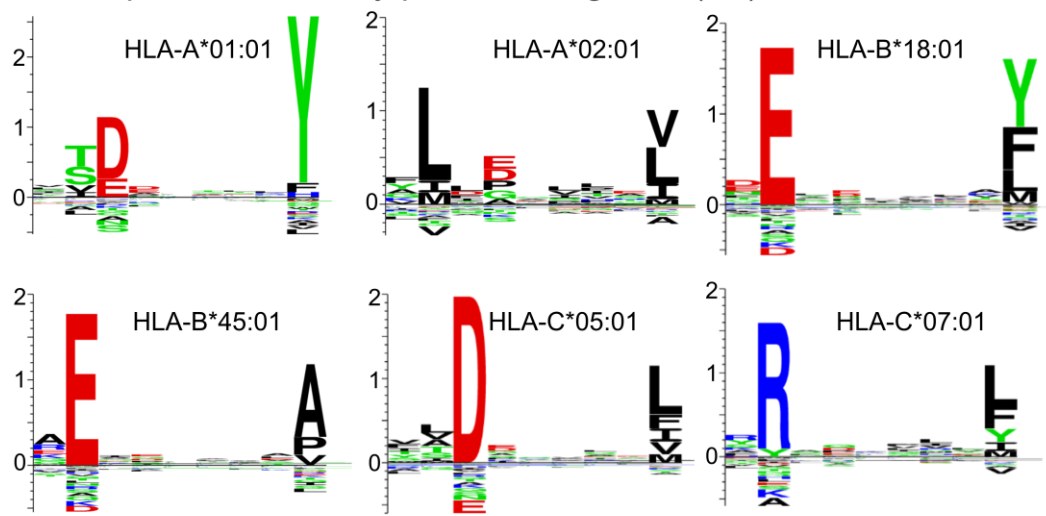

**Supplementary Figure S5. GibbsCluster 2.0 of HCT-116 label-free identified peptides.** (A) Identified peptides of length 8 to 12 were subjected to unsupervised alignment and clustering by GibbsCluster 2.0<sup>6</sup> using recommended settings. For each cluster, sequence logos were made using Logomaker<sup>7</sup> (*Left*) and the distribution of NetMHCpan-4.1<sup>5</sup> predicted binding level was displayed (*Middle*). For predicted weak and strong binders the HLA allele distribution was displayed (*Right*). (B) NetMHCpan-4.1 motif viewer of HCT-116 alleles (naturally presented ligands, EL). Abbreviations: NB, non-binder; SB, strong binder; WB, weak binder.

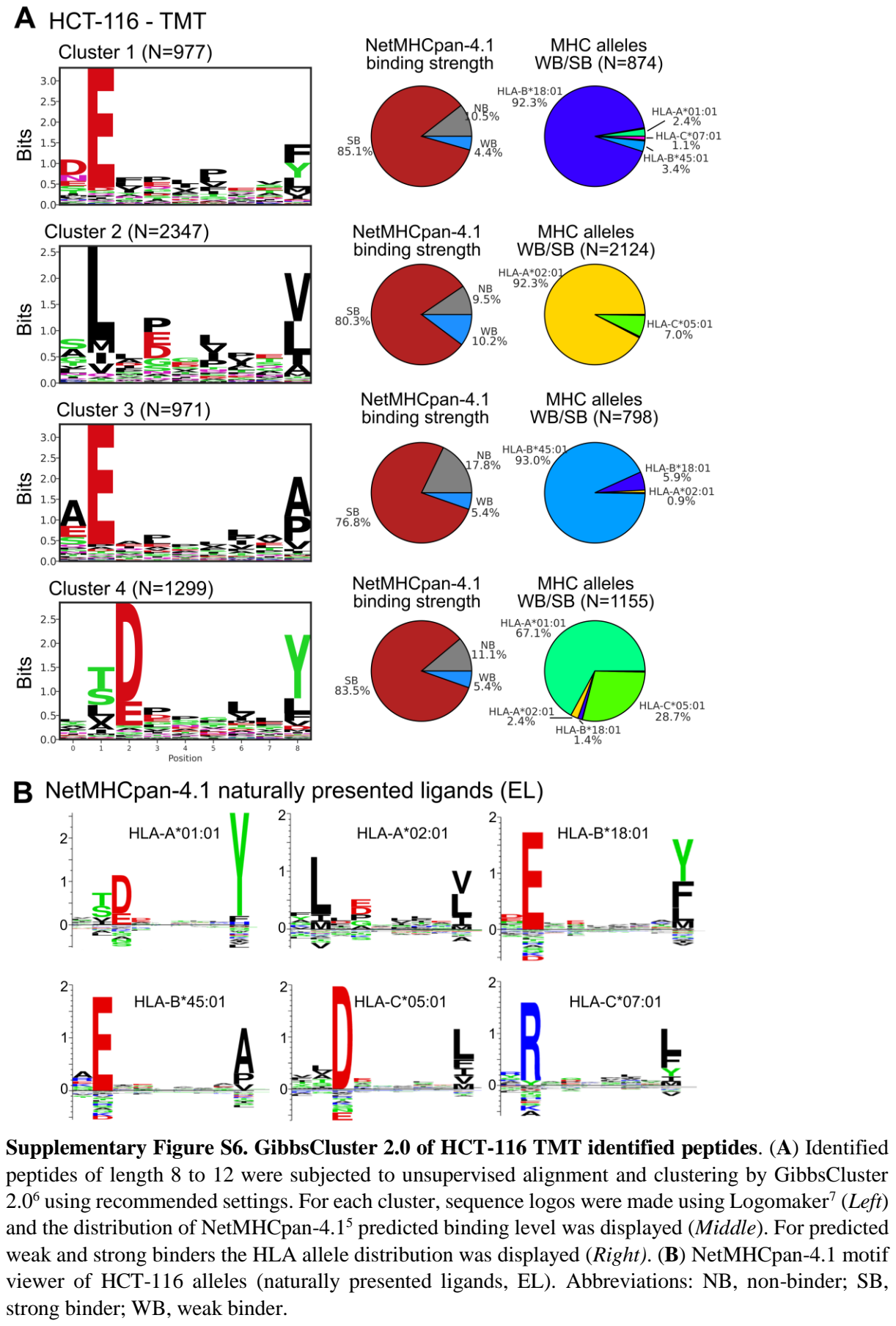

### A HCT-116 - label free

Strong binder HLA-B\*18:01 (N=641)

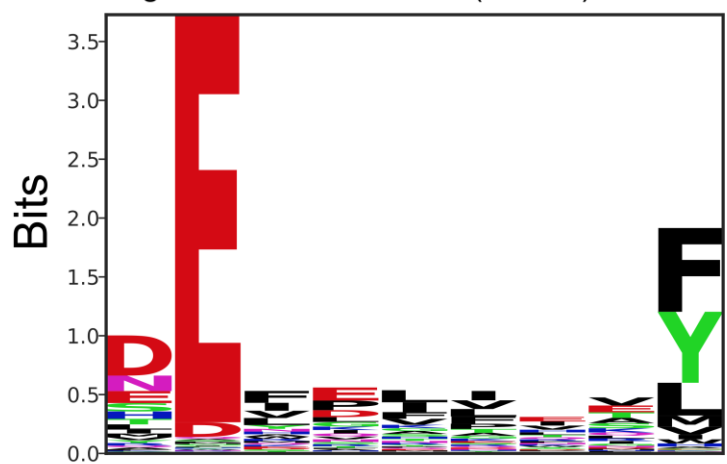

Peptide length histogram

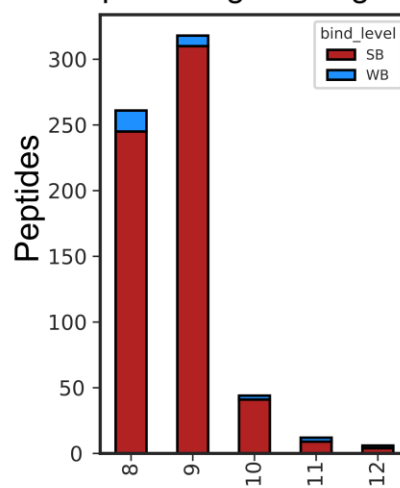

### B HCT-116 - TMT

Strong binder HLA-B\*18:01 (N=873)

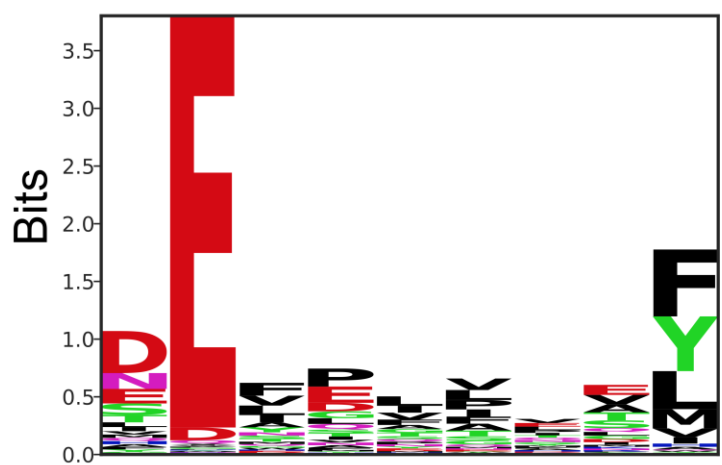

Peptide length histogram

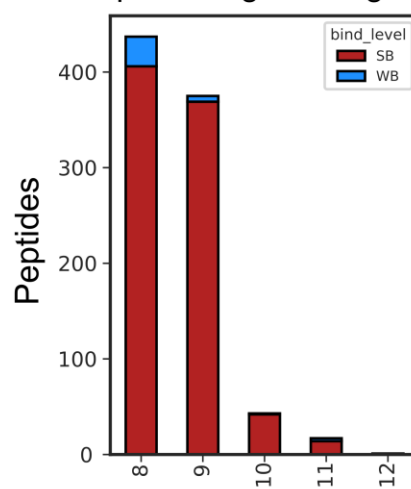

**Supplementary Figure S7. HLA-B\*18:01 shows binding preference to 8- and 9-mers.** Sequence logo of peptides predicted as weak binder (WB, %rank < 2) or strong binder (SB, %rank < 0.5) to HLA-B\*18:01 by NetMHCpan-4.1<sup>5</sup>. Only peptides for which HLA-B\*18:01 was predicted as the best HLA binder were included. Logos were made by Logomaker<sup>7</sup> using the predicted peptide binding core (*Left*). A peptide length histogram was plotted for WB and SBs (*Right*).

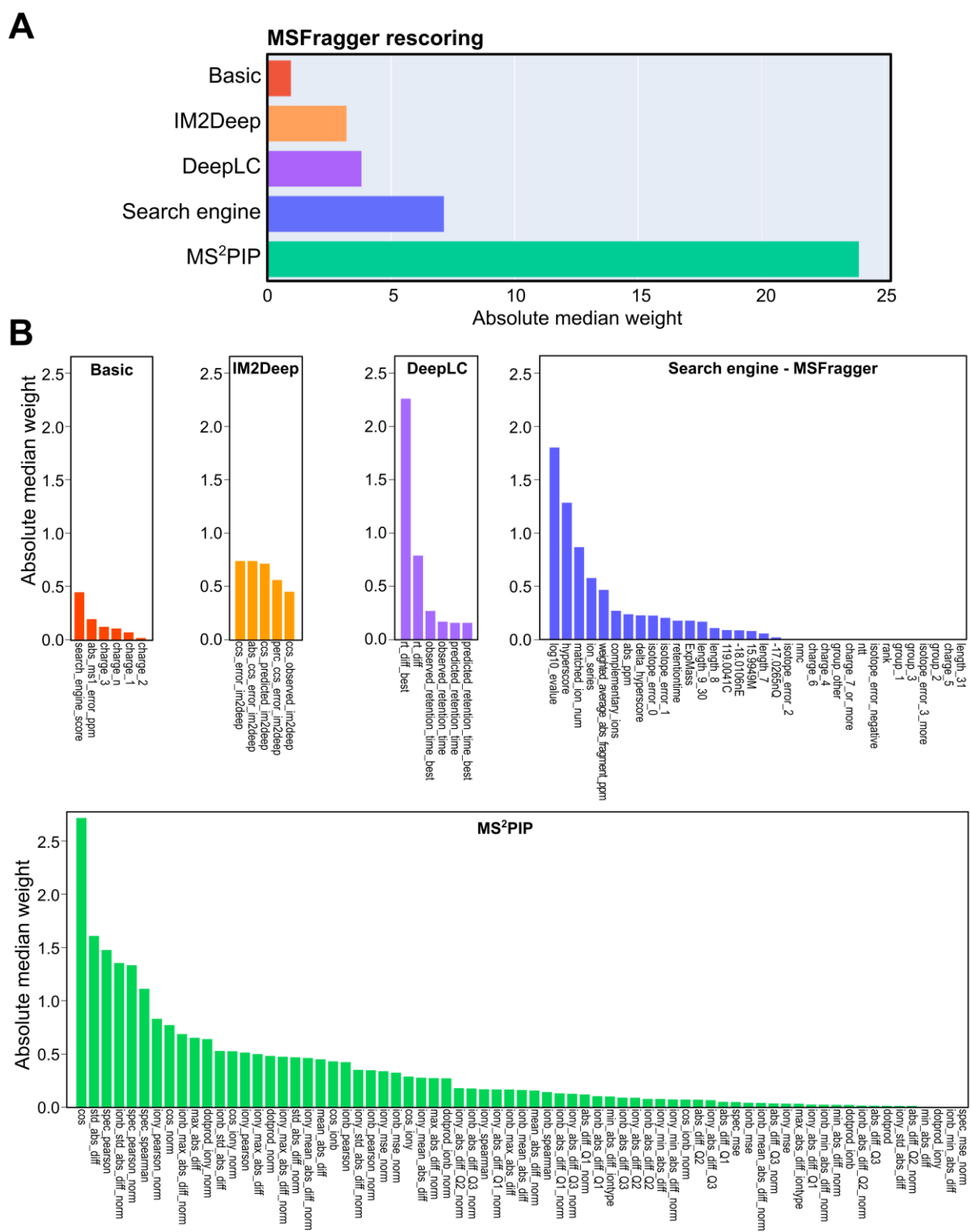

**Supplementary Figure S8. TIMS<sup>2</sup>Rescore feature weights in MSFragger rescoring for MHC class I immunopeptides in HeLa label-free samples. (A)** Absolute median weight of features summed per feature generator, either basic features (red), IM2Deep<sup>8</sup> for collisional-cross section (CCS) rescoring (orange), DeepLC<sup>9</sup> (purple) for retention time rescoring, MSFragger<sup>10</sup> search engine score rescoring (blue), and MS<sup>2</sup>PIP<sup>11</sup> for fragment ion intensity rescoring. **(B)** Absolute median weight per feature colored per feature generator. Weights were extracted from TIMS<sup>2</sup>Rescore<sup>8</sup> generated reports, which are deposited for every rescoring at deposited at the Open Science Framework (OSF) project with DOI 10.17605/OSF.IO/7DKSA.

**A**

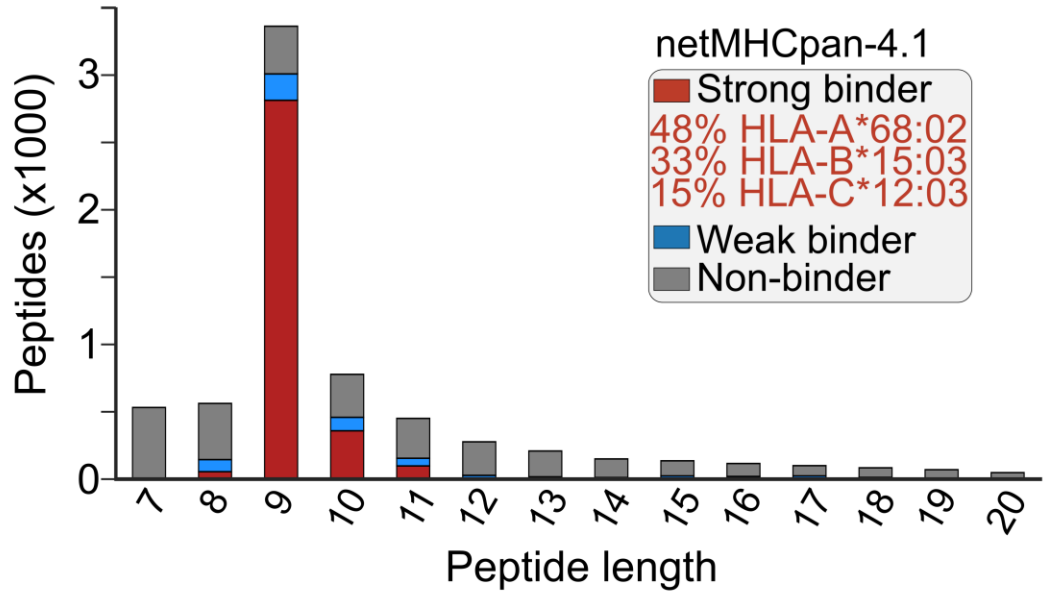

**B**

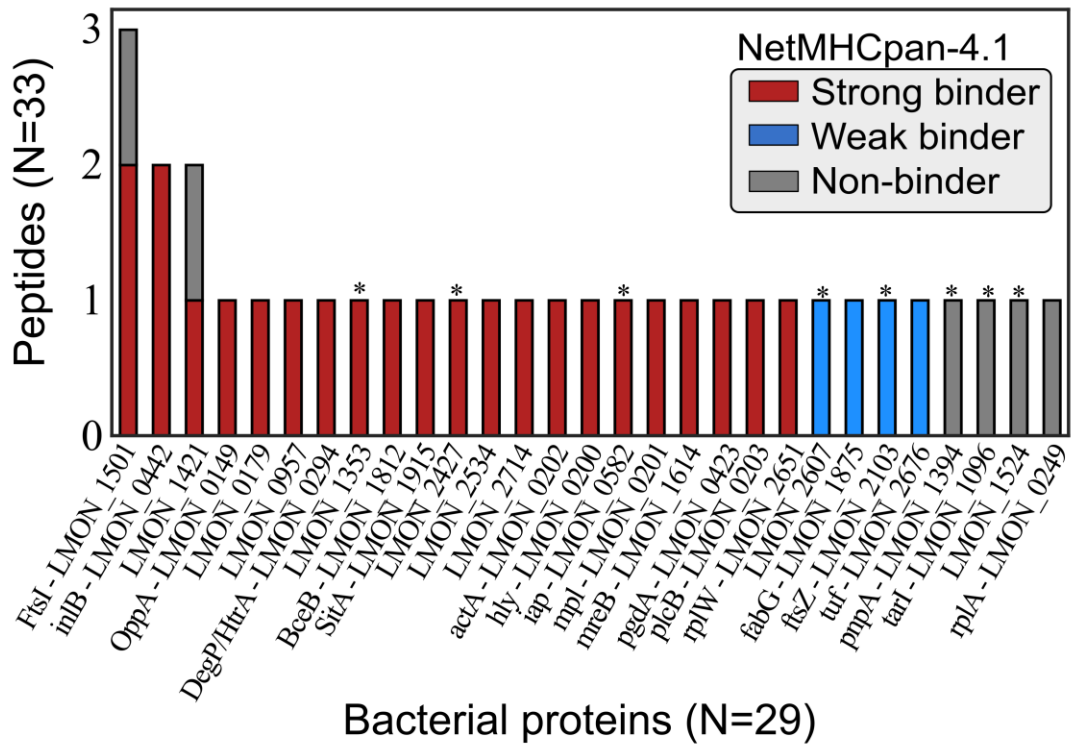

**Supplementary Figure S9. TimsTOF SCP identifies novel antigens in *Listeria*-infected HeLa samples.** (A) The number of unique peptide sequences identified per amino acid length. Peptide sequences predicted as strong binder (SB, %rank < 0.5) or weak binder (WB, %rank < 2) by NetMHCpan-4.1<sup>5</sup> were indicated in red and blue, respectively. Other peptides (non-binder, NB) were indicated in grey. The proportion of major HLA alleles (>10%, best %rank per peptide) is indicated by red text for SB peptides. (B) Bacterial protein histogram, displaying the number of identified bacterial peptides per protein. Immunoepitopes with NetMHCpan-4.1 predicted SB, WB and NB were indicated in red, blue and grey, respectively. Bacterial proteins not yet identified in the Q Exactive HF (re-)analysis were marked by an asterisk (\*).

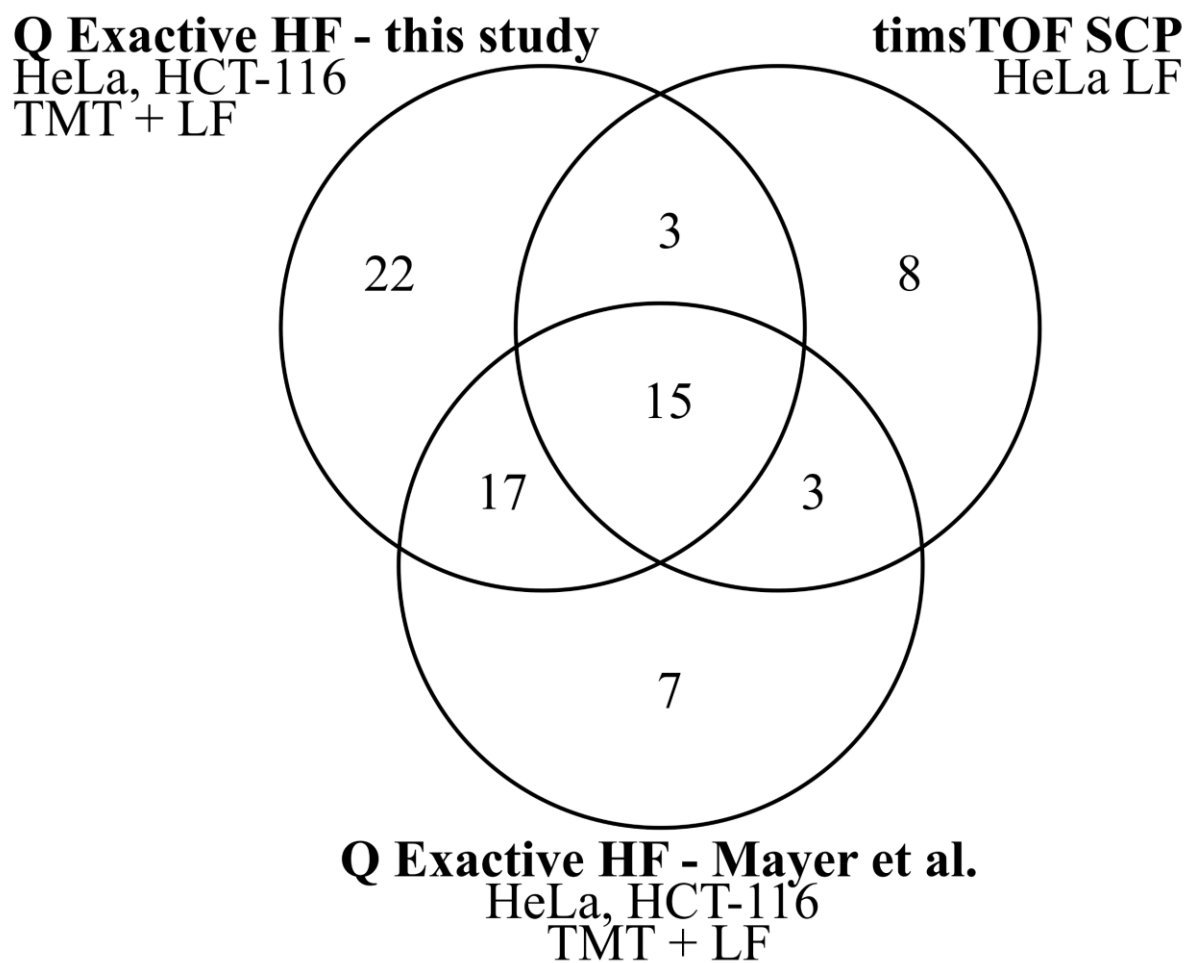

**Supplementary Figure S10. Novel *Listeria* proteins identified in this study.** Venn diagram of *Listeria* proteins identified in this study compared to Mayer et al.<sup>4</sup>, with a total of 33 novel identified proteins. Note that timsTOF SCP data was only generated for HeLa label-free (LF) data.

**Supplementary Table 1. Set polygon vertex points for timsTOF SCP re-injection of *Listeria*-infected HeLa label-free samples.**

| IMS polygon filter mass | IMS polygon filter mobility |
| --- | --- |
| 270.54 | 0.551 |
| 400.74 | 0.850 |
| 702.59 | 1.101 |
| 704.02 | 1.723 |
| 1716 | 1.764 |

**Supplementary Table 2. Set collision energy scheme for timsTOF SCP re-injection of *Listeria*-infected HeLa label-free samples.**

| 1/K0 [Vs cm<sup>-2</sup>] | Collision energy [eV] |
| --- | --- |
| 0.7 | 20 |
| 1.06 | 30 |
| 1.16 | 40 |
| 1.34 | 40 |
| 1.68 | 70 |
